# Improved phylogenetic resolution within Siphonophora (Cnidaria) with implications for trait evolution

**DOI:** 10.1101/251116

**Authors:** Catriona Munro, Stefan Siebert, Felipe Zapata, Mark Howison, Alejandro Damian Serrano, Samuel H. Church, Freya E. Goetz, Philip R. Pugh, Steven H.D. Haddock, Casey W. Dunn

**Author notes:** Authors contributed equally (Catriona Munro, Stefan Siebert).

## Abstract

Siphonophores are a diverse group of hydrozoans (Cnidaria) that are found at all depths of the ocean - from the surface, like the familiar Portuguese man of war, to the deep sea. Siphonophores play an important role in ocean ecosystems, and are among the most abundant gelatinous predators. A previous phylogenetic study based on two ribosomal RNA genes provided insight into the internal relationships between major siphonophore groups, however there was little support for many deep relationships within the clade Codonophora. Here, we present a new siphonophore phylogeny based on new transcriptome data from 30 siphonophore species analyzed in combination with 13 publicly available genomic and transcriptomic datasets. We use this new phylogeny to reconstruct several traits that are central to siphonophore biology, including sexual system (monoecy vs. dioecy), gain and loss of zooid types, life history traits, and habitat. The phylogenetic relationships in this study are largely consistent with the previous phylogeny, but we find strong support for new clades within Codonophora that were previously unresolved. These results have important implications for trait evolution within Siphonophora, including favoring the hypothesis that monoecy arose twice.

## 1. Introduction

Siphonophores (Fig. 1 and 2) are among the most abundant gelatinous predators in the open ocean, and have a large impact on ocean ecosystems (Choy et al., 2017; Pagès et al., 2001; Pugh, 1984; Pugh et al., 1997; Purcell, 1981; Williams and Conway, 1981). Siphonophores, which belong to Hydrozoa (Cnidaria), are found at all depths in the ocean. The most familiar species is the Portuguese man of war *Physalia physalis*, which floats at the surface and can wash up conspicuously onto beaches (Totton, 1960). Most species are planktonic, living in the water column, where some grow to be more than 30 meters in length (Mackie et al., 1987). There is also a small clade of benthic siphonophores, Rhodaliidae (Pugh, 1983), that are tethered to the bottom for part of their lives. There are currently 187 valid described siphonophore species.

**Figure 1:**
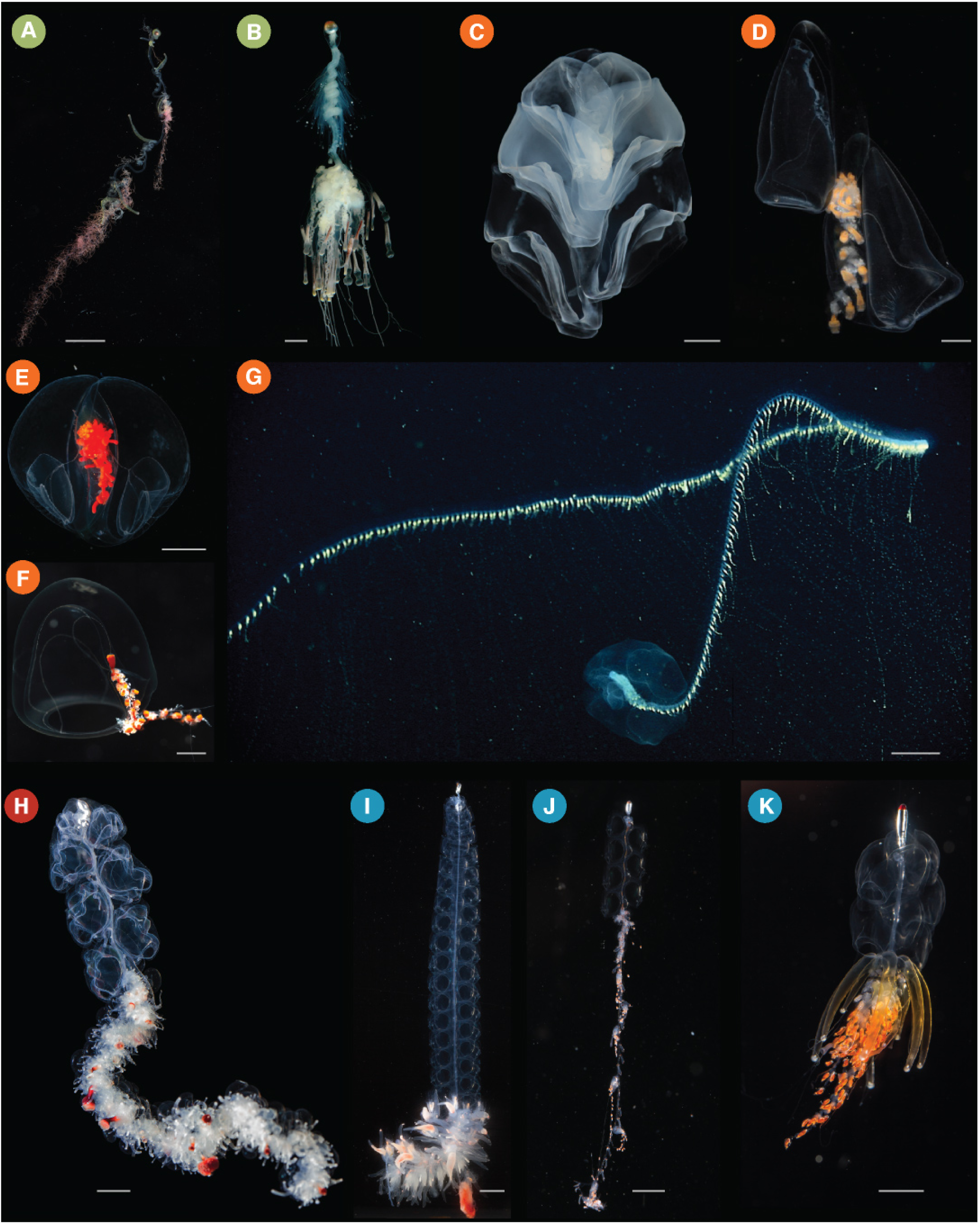
Photographs of living siphonophores. They correspond to the clades shown in Figure 3 as follows: Cystonectae (A-B), Calycophorae (C-G), Apolemiidae (H), and Clade A within Euphysonectae (I-K). (A) *Rhizophysa eysenhardtii*, scale bar = 1 cm. (B) *Bathyphysa conifera*, scale bar = 2cm. (C) *Hippopodius hippopus*, scale bar = 5 mm. (D) *Kephyes hiulcus*, scale bar = 2 mm. (E) *Desmophyes haematogaster*, scale bar = 5 mm. (F) *Sphaeronectes christiansonae*, scale bar = 2 mm. (G) *Praya dubia*, scale bar = 4 cm. (H) *Apolemia* sp., scale bar = 1 cm. (I) *Lychnagalma utricularia*, scale bar = 1 cm. (J) *Nanomia* sp., scale bar = 1 cm. (K) *Physophora hydrostatica*, scale bar = 5 3mm. Photo credits: S. Siebert (C,H,I,K), S. Haddock (A,D,E,F), R. Sherlock (B), MBARI (G), C. Munro (J)

**Figure 2:**
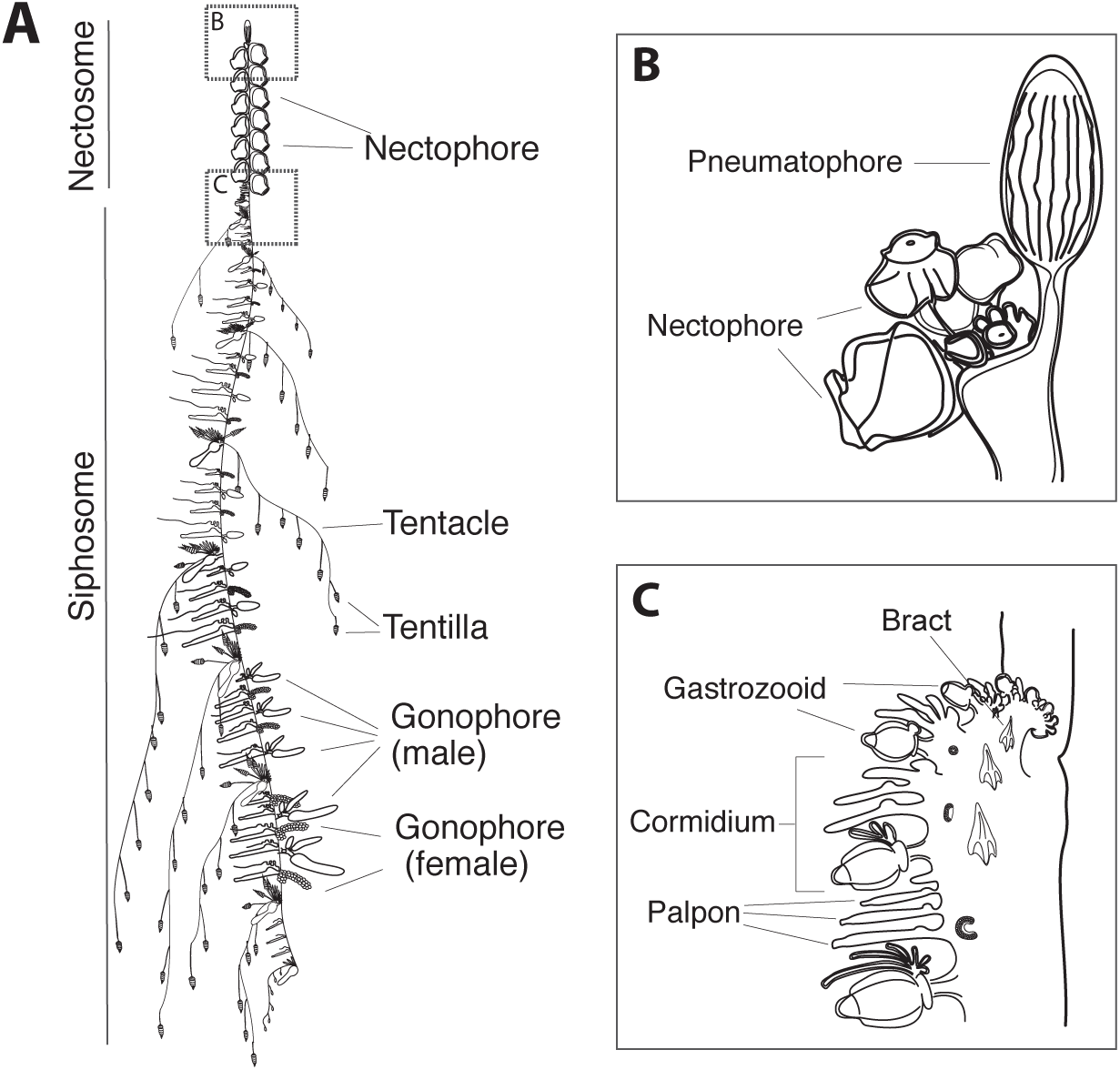
Schematic of the siphonophore *Nanomia bijuga*, orientated with the anterior of the colony at the top of the page, and the ventral side to the left. Adapted from http://commons.wikimedia.org/wiki/File:Nanomia_bijuga_whole_animal_and_growth_zones.svg, drawn by Freya Goetz. (A) Overview of the whole mature colony. (B) Inset of the pneumatophore and nectosomal growth zone. A series of buds give rise to nectophores. (C) Inset of the siphosomal growth zone. Probuds subdivide to give rise to zooids in repeating-units (cormidia). The gastrozooid (specialized feeding polyp) is the posterior-most zooid within each cormidium.

Siphonophores remain poorly known, in large part because they are fragile and difficult to collect. They have, however, been of great interest for more than 150 years due to their unique structure and development (Mackie et al., 1987; Mapstone, 2014). Like many other cnidarians, they are colonial: they grow by incomplete asexual reproduction. Each colony arises from a single embryo that forms the protozooid, the first body. One or two growth zones (Fig. 2) then arise that asexually produce other genetically identical zooids that remain attached (Carré and Carré, 1995, Carré 1991; Carré, 1969, Carré 1967). These zooids are each homologous to a solitary animal, but are physiologically integrated (Dunn and Wagner, 2006; Mackie et al., 1987; Totton, 1965). Siphonophores differ significantly from other colonial animals in terms of colony structure and development – their zooids are highly functionally specialized and arranged in precise, repeating, species-specific patterns (Beklemishev, 1969; Cartwright and Nawrocki, 2010). The functions that zooids are specialized for include feeding, reproducing, or swimming (Fig. 2) (Dunn and Wagner, 2006).

Understanding the unique ecology, morphology, and development of siphonophores requires a well-resolved phylogeny of the group. The relationship of siphonophores to other hydrozoans has been difficult to resolve (Cartwright et al., 2008; Cartwright and Nawrocki, 2010; Zapata et al., 2015), but there has been progress on their internal relationships. A phylogeny (Dunn et al., 2005) based on two genes (16S, 18S) from 52 siphonophore taxa addressed several long standing questions about siphonophore biology. Including the relationships of the three historically recognised groups, Cystonectae, Physonectae, and Calycophorae. Cystonectae was found to be sister to all other siphonophores, while Calycophorae were nested within “Physonectae”. The name Codonophora was given to this clade of “Physonectae” and Calycophorae (Dunn et al., 2005).

Major questions remained after this early work, though. There was, in particular, little support for important deep relationships within Codonophora. Understanding these relationships is key to resolving the evolution of several traits of importance, including sexual systems (monoecy versus dioecy) and the gain and loss of particular zooids, such as palpons (Fig. 2). Here we present a broadly sampled phylogenetic analysis of Siphonophora that considers transcriptomic data from 33 siphonophore species and 10 outgroup species (2 outgroups were subsequently excluded due to poor sampling). Using 1,071 genes, we find strong support for many relationships found in the earlier phylogeny (Dunn et al., 2005), and also provide new resolution for key relationships that were unresolved in that previous study. Using this phylogeny, we reconstruct the evolutionary history of characters central to the unique biology of siphonophores, including zooid type, life history traits, and habitat.

## 2. Material and methods

All scripts for the analyses are available in a git repository at https://github.com/caseywdunn/siphonophore_phylogeny_2017. The most recent commit at the time of the analysis presented here was 7c33159bed8a993b09bb1908cf9bd317c04ef3d2.

### 2.1 Collecting

Specimens were collected in the north-eastern Pacific Ocean, Mediterranean, and the Gulf of California. Collection data on all examined specimens, a description of the tissue that was sampled from the colony, collection mode, sample processing details, mRNA extraction methods, sequencing library preparation methods, and sequencing details are summarized in the file Supplementary data 1 (also found in the git repository). Monterey Bay and Gulf of California specimens were collected by remotely operated underwater vehicle (ROV) or during blue-water SCUBA dives. *Chelophyes appendiculata* and *Hippopodius hippopus* (Fig. 1C) specimens were collected in the bay of Villefranche-sur-Mer, France, during a plankton trawl on 13 April 2011. Available physical vouchers have been deposited at the Museum of Comparative Zoology (Harvard University), Cambridge, MA, the Peabody Museum of Natural History (Yale University), New Haven, CT, or had been previously deposited at the United States National Museum (Smithsonian Institution), Washington, DC. Accession numbers are given in Supplementary data 1. In cases where physical vouchers were unavailable we provide photographs to document species identity (see git repository).

### 2.2 Sequencing

When possible, specimens were starved overnight in filtered seawater at temperatures close to ambient water temperatures at the time of specimen collection. mRNA was extracted directly from tissue using a variety of methods (Supplementary data 1): Magnetic mRNA Isolation Kit (NEB, #S1550S), Invitrogen Dynabeads mRNA Direct Kit (Ambion, #61011), Zymo Quick RNA MicroPrep (Zymo #R1050), or from total RNA after Trizol (Ambion, #15596026) extraction and through purification using Dynabeads mRNA Purification Kit (Ambion, #61006). In case of anticipated very small total RNA quantities, only a single round of bead purification was performed. Extractions were performed according to the manufacturer’s instruction. All samples were DNase treated (TURBO DNA-free, Invitrogen #AM1907; or on column DNase treatment with Zymo Quick RNA MicroPrep). Libraries were prepared for sequencing using the Illumina TruSeq RNA Sample Prep Kit (Illumina, #FC-122-1001, #FC-122-1002), the Illumina TruSeq Stranded Library Prep Kit (Illumina, #RS-122-2101) or the NEBNext RNA Sample Prep Master Mix Set (NEB, #E6110S). We collected long read paired end Illumina data for *de novo* transcriptome assembly. In the case of large tissue inputs, libraries were sequenced separately for each tissue, subsequently subsampled and pooled *in silico*. Libraries were sequenced on the HiSeq 2000, 2500, and 3000 sequencing platforms. Summary statistics for each library are given in the file Supplementary data 2. All sequence data have been deposited in the NCBI sequence read archive (SRA) with Bioproject accession number PRJNA255132.

### 2.3 Analysis

New data were analysed in conjunction with 13 publicly available datasets, with a total number of 43 species. Sequence assembly, annotation, homology evaluation, gene tree construction, parsing of genes trees to isolate orthologous sequences, and supermatrix construction were conducted with Agalma v. 1.0.0 (Dunn et al., 2013; Guang et al., 2017). This workflow integrates a variety of existing tools (Altschul et al., 1990; Enright et al., 2002; Grabherr et al., 2011; Katoh and Standley, 2013; Langmead and Salzberg, 2012; Li and Dewey, 2011; Li et al., 2009; Sukumaran and Holder, 2010; Talavera and Castresana, 2007) and new methods. Maximum likelihood analyses of the supermatrix were conducted with RAxML v 8.2.0 (Stamatakis, 2006) and implemented via Agalma. Bayesian Inference (BI) analyses of the supermatrix were conducted using Phylobayes v. 1.7a-mpi (Lartillot et al., 2009). Sequence alignments, sampled and consensus trees, and voucher information are available in the git repository. Tree figures were rendered with ggtree (Yu et al., 2016).

Two outgroup species, *Atolla vanhoeffeni* and *Aegina citrea*, were removed from the final supermatrix due to low gene occupancy (gene sampling of 20.8%sss and 14.5%sss respectively in a 50%sss occupancy matrix with 2,203 genes). The final analyses presented here consider 33 siphonophore species and 8 outgroup species. This includes new data for 30 species. In the final analyses, we sampled 1,071 genes to generate a supermatrix with 60%sss occupancy and a length of 378,468 amino acids (Fig. S1).

ML analyses were conducted on the unpartitioned supermatrix using the WAG+Γ model of amino acid substitution, and bootstrap values were estimated using 1000 replicates. BI was conducted using two different CAT models, CAT-Poisson and CAT-GTR (Lartillot and Philippe, 2004). Two independent MCMC chains were run under the CAT-GTR model, and four independent MCMC chains were run under the CAT-Poisson model. The BI analyses did not converge (CAT-Poisson: maxdiff=1, meandiff=0.0148; CAT-GTR: maxdiff=1, meandiff=0.0127). Only the results from the CAT-Poisson model are presented here. Visual inspection of the traces indicated that a burn in of 400 trees was sufficient for all CAT-Poisson runs. This left 15847 trees in the posterior.

We used the Swofford-Olsen-Waddell-Hillis (SOWH) test (Swofford et al., 1996) to evaluate two hypotheses (Fig. 3C, S2): (i) “Physonectae” is monophyletic (Totton, 1965); (ii) monoecious species are monophyletic (Dunn et al., 2005). As the sexual system of *Rudjakovia* sp. is unclear, we carried out two tests of the monophyly of monoecy, one with *Rudjakovia* sp. included as a monoecious species, and one without (Fig. 3C, S2). We used SOWHAT (Samuel H. Church et al., 2015) dev. version 0.39 (commit fd68ef57) to carry out the SOWH tests in parallel with the default options and an initial sample size of 100 (analysis code can be found in the git repository). For each hypothesis we defined a topology with a single constrained node that was inconsistent with the most likely topology (Fig. 3). We used a threshold for significance of 0.05 and following the initial 100 samples, we evaluated the confidence interval around the p-value to determine if more samples were necessary.

**Figure 3:**
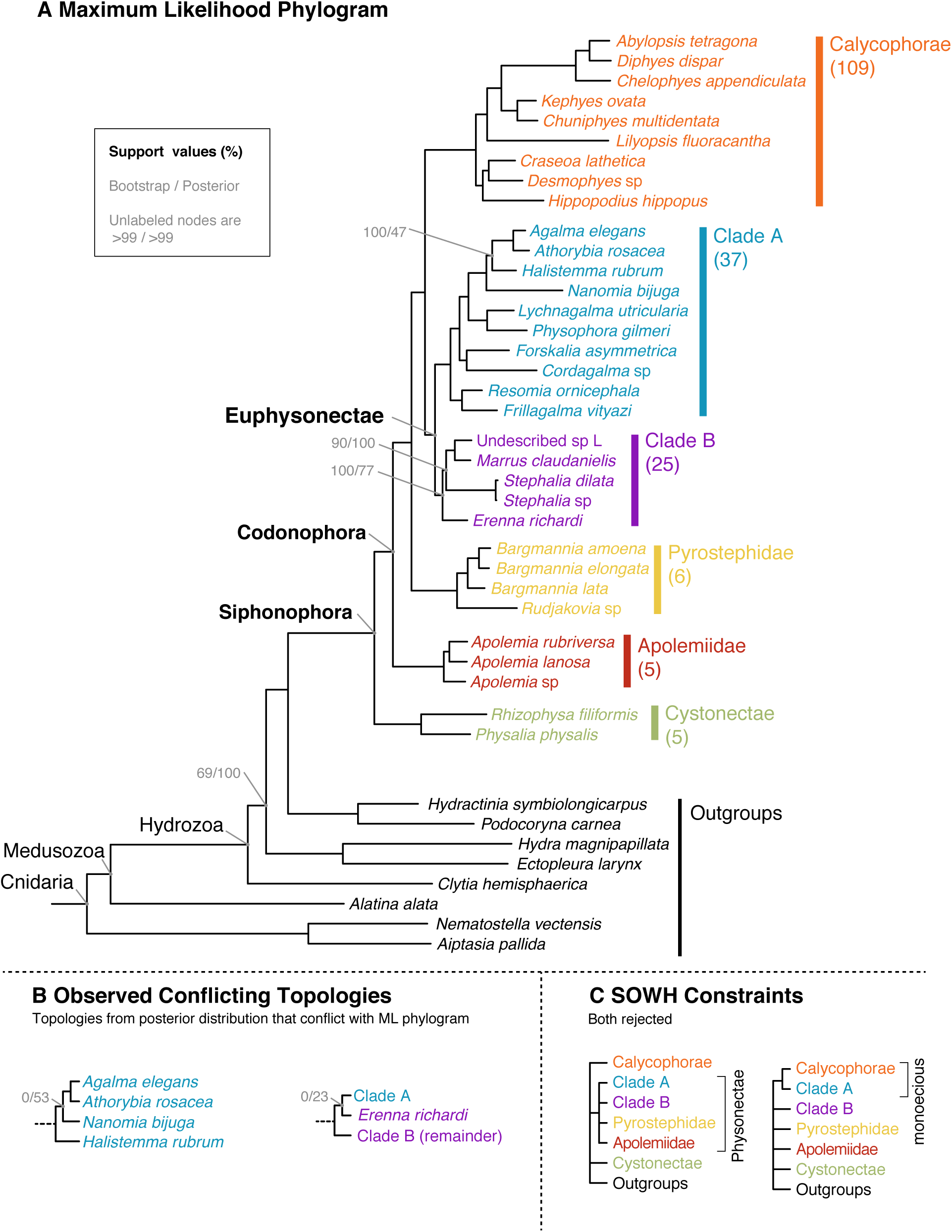
(A) Maximum likelihood (ML) phylogram with bipartition frequencies from the ML bootstraps and the Bayesian posterior distribution of trees. Unlabeled nodes have support >0.99 for both bootstraps and posteriors. The numbers of valid described species estimated to be based in each clade based on taxonomy are shown below each clade name on the right. (B) The topologies found in the posterior distribution of trees that conflict with the ML tree. (C) The topologies evaluated by the SOWH tests. For more details on the SOWH topologies refer to Fig. S2.

Morphological character data used in trait mapping were obtained from the literature or direct observation of available voucher material. Depth distribution data was queried from the MBARI VARS database (http://www.mbari.org/products/research-software/video-annotation-and-reference-system-vars/) (Schlining and Stout, 2006). We used stochastic character mapping to infer the probable evolution of traits on the tree in R using the phytools package (Huelsenbeck et al., 2003; Revell, 2012). Subsequent analyses were conducted in R and integrated into this manuscript with the knitr package. See Supplementary Information for R package version numbers.

## 3. Results and Discussion

### 3.1 Species phylogeny and hypothesis testing

Most relationships received strong support across analysis methods (Fig. 3A), with a couple of localized exceptions (Fig. 3B). The phylogenetic relationships recovered in this study are largely consistent with those found in a previous study based on two genes (16S and 18S ribosomal RNA) (Dunn et al., 2005). Relationships that receive strong support in both studies include the placement of Cystonectae as sister to Codonophora (the clade that includes all other siphonophores), the placement of Apolemiidae as sister to all other codonophorans, and the placement of Calycophorae within the paraphyletic “Physonectae”. Multiple nodes that were not resolved in the previous two-gene analysis receive strong support in the present 1,071-gene transcriptome analyses. There is strong support for Pyrostephidae as sister to all other non-apolemiid codonophorans. Within the clade that is sister to Pyrostephidae, we find two main clades, Calycophorae and a clade we here name Euphysonectae (Fig. 3A). It includes the remaining non-apolemiid, non-pyrostephid “Physonectae”. We define Euphysonectae as the clade consisting of *Agalma elegans* and all taxa that are more closely related to it than to *Diphyes dispar*.

Euphysonectae consists of two reciprocally monophyletic groups that we here provisionally refer to as Clade A and Clade B (Fig. 3A). The presence of an involucrum, a fold around the base of the cnidoband (Totton, 1965), is a potential synapomorphy for Clade A. Species of Clade A also have a descending mantle canal within the nectophores (Fig. S6), a structure that is also present in some calycophorans. Members of Clade A are also monoecious (Fig. 5). There is not a clear synapomorphy for Clade B. Within Clade B there is low support for the placement of *Erenna richardi*, which is placed as sister to Clade A in some analyses (Fig. 3B). More taxon sampling will be required to determine the relationship of species within this clade.

Within Clade A, *Physophora gilmeri* along with *Lychnagalma utricularia* (Fig. 1I) (both not included in the previous phylogeny) are sister to Agalmatidae, a clade restricted to *Agalma*, *Athorybia*, *Melophysa*, *Halistemma* and *Nanomia* (Dunn et al., 2005; Pugh, 2006). In the rDNA study, *P. hydrostatica* (the presumed sister species to *P. gilmeri*) was sister to Forskaliidae with low support. The position of *Cordagalma cordiforme* (= *Cordagalma ordinatum*) (Pugh, 2016) was previously unresolved, while in this analysis *Cordagalma* sp. is in a clade with *Forskalia asymmetrica*, falling outside of Agalmatidae. Placement of *Cordagalma* outside Agalmatidae is consistent with previous analyses of morphological and molecular data (Dunn et al., 2005; Pugh, 2006).

Within Calycophorae, taxon sampling is shallower here than in the previous study. The calycophoran relationships that can be investigated, however, are in broad agreement with the previous analysis. Calycophorans have in the past been split into two groups, prayomorphs and diphyomorphs, based on morphology after Mackie et al. (1987). As in the previous study, the results presented here indicate that the prayomorphs are paraphyletic with respect to the diphyomorphs. *Craseoa lathetica* and *Desmophyes* sp. are sister to *Hippopodius hippopus* in this study, while in the previous study, the relationship between *C. lathetica* and the clade including *H. hippopus* was unresolved.

**Table 1:** A complete list of specimens collected for this work. New data indicated by Y, blank fields indicate that data were already published.

| New data | Species | SRA Number | Depth (m) | Lat Lon |
| --- | --- | --- | --- | --- |
|  | Agalma elegans |  | 3–20 | 35.56 N 122.55 W |
| Y | Bargmannia elongata | SRR1548343;SRR1548344;SRR1548345;SRR1548346;SRR1548347 | 412/805/636/818 | 36.12 N 122.67 W |
| Y | Frillagalma vityazi | SRR1548362;SRR1548363;SRR1548364 | 407 | 36.69 N 122.05 W |
| Y&N | Nanomia bijuga | SRR1548376;SRR1548377;SRR871527 | 414/387 | 36.60 N 122.15 W |
|  | Physalia physalis | SRR871528 |  |  |
|  | Abylopsis tetragona | SRR871525 |  |  |
|  | Aegina citrea | SRS893439 |  |  |
|  | Aiptasia pallida | SRR6967; SRR6967; SRR6967 |  |  |
|  | Alatina alata | SRR1952741 |  |  |
| Y | Apolemia rubriversa | SRR1548342 | 767 | 36.70 N 122.05 W |
|  | Atolla vanhoeffeni | SRR1952729 |  |  |
| Y | Chelophyes appendiculata | SRR1548354 | 3–20 |  |
| Y | Chuniphyes multidentata | SRR1548355 | 327 | 36.79 N 122.00 W |
|  | Clytia hemisphaerica |  |  |  |
| Y | Cordagalma sp | SRR1548356 | 252 | 36.70 N 122.06 W |
|  | Ectopleura larynx | SRR923510 |  |  |
| Y | Erenna richardi | SRR1548360 | 1044 | 36.61 N 122.38 W |
| Y | Forskalia asymmetrica | SRR1548361 | 253 | 36.80 N 122.00 W |
| Y | Hippopodius hippopus | SRR1548371 | 3–20 | 43.69 N 7.315 E |
|  | Hydra magnipapillata |  |  |  |
|  | Hydractinia symbiolongicarpus | SRX474878 |  |  |
| Y | Kephyes ovata | SRR1548372 | 452 | 36.36 N 122.81 W |
| Y | Lilyopsis fluoracantha | SRR1548373 | 320 | 36.69 N 122.04 W |
| Y | Lychnagalma utricularia | SRR1548374 | 431 | 36.69 N 122.04 W |
| Y | Marrus claudanielis | SRR1548375 | 1427 | 36.07 N 122.29 W |
|  | Nematostella vectensis |  |  |  |
| Y | Undescribed Sp. L | SRR1548381 | 1463 | 36.70 N 122.57 W |
| Y | Podocoryna carnea | SRR1266262 |  |  |
| Y | Desmophyes sp. | SRR1548358 | 1363 | 35.48 N 123.64 W |
|  | Craseoa lathetica | SRR871529 |  |  |
| Y | Resomia ornicephala | SRR1548382 | 322 | 35.48 N 123.86 W |
| Y | Rhizophysa filiformis | SRR1548383 | 10 | 27.23 N 110.46 W |
| Y | Stephalia dilata | SRR1548384 | 3074 | 35.62 N 122.67 W |
| Y | Apolemia lanosa |  | 1073 | 36.70 N 122.08 W |
| Y | Apolemia sp |  | 461 | 36.60 N 122.15 W |
| Y | Bargmannia amoena |  | 1251 | 36.70 N 122.08 W |
| Y | Bargmannia lata |  | 1158 | 36.067 N 122.30 W |
| Y | Rudjakovia sp |  | 334 | 36.00 N 122.42 W |
| Y | Stephalia sp |  | 3255 | 36.39 N 122.67 W |
| Y | Physophora gilmeri |  | 242 | 36.36 N 122.40 W |
| Y | Halistemma rubrum |  | 313 | 24.68 N 109.90W |
| Y | Athorybia rosacea |  | 3–20 | 22.92 N 108.36 W |
| Y | Diphyes dispar |  | 3–20 | 35.93 N 122.93 W |

Using the Swofford-Olsen-Waddell-Hillis (SOWH) test (Swofford et al., 1996), we evaluated the following three alternative phylogenetic hypotheses against the most likely tree topology (Fig. 3C): (i) monophyletic Physonectae, (ii) and (iii) monophyletic monoecious siphonophores, with and without *Rudjakovia* sp. respectively. In all three tests the alternative hypothesis was rejected (p-value <0.01, confidence interval: <0.001 − 0.03, Fig. S2).

The broad sampling approach of this phylogeny provides new evidence for the relationships between major siphonophore clades within Codonophora, specifically between Pyrostephidae, Calycophorae, and the newly named Euphysonectae. This opens up new questions about key relationships within both Calycophorae and Euphysonectae – where future transcriptome sampling efforts should be focused. Within Euphysonectae, two clades (Clade A and Clade B) are hypothesized, however there is weaker support for Clade B (Fig. 3A, 3B). Expanding sampling of species that probably fall in Clade B, including other *Erenna* species, rhodaliids, and relatives of Undescribed sp L, will greatly expand our understanding of these two groups and perhaps provide evidence of Clade B synapomorphies. Similarly, within Calycophorae, increased taxon sampling is needed. This study, and the previous phylogenetic study (Dunn et al., 2005), suggest that the prayomorphs are paraphyletic, but for slightly different reasons given the different sampling of the analyses. In Dunn et al. (2005), a clade of prayomorphs including *Praya dubia* (Fig. 1G), *Nectadamas diomedeae*, and *Nectopyramis natans* (not included in this study) were found to be sister to all other calycophorans, while in this study, the prayomorph *Lilyopsis fluoracantha* (not included in the previous study) is found in a clade including diphyomorph calycophorans that is sister to all other prayomorphs. Expanded taxon sampling, particularly *P. dubia* or a nectopyramid, but also extensive sampling across the major prayomorph and diphyomorph groups, will expand our understanding of relationships within Calycophorae. This will be especially important for understanding trait evolution within Calycophorae, for example, the release of eudoxids (Fig. 4), or the arrangement of male and female zooids along the stem (see section 3.2 below).

**Figure 4:**
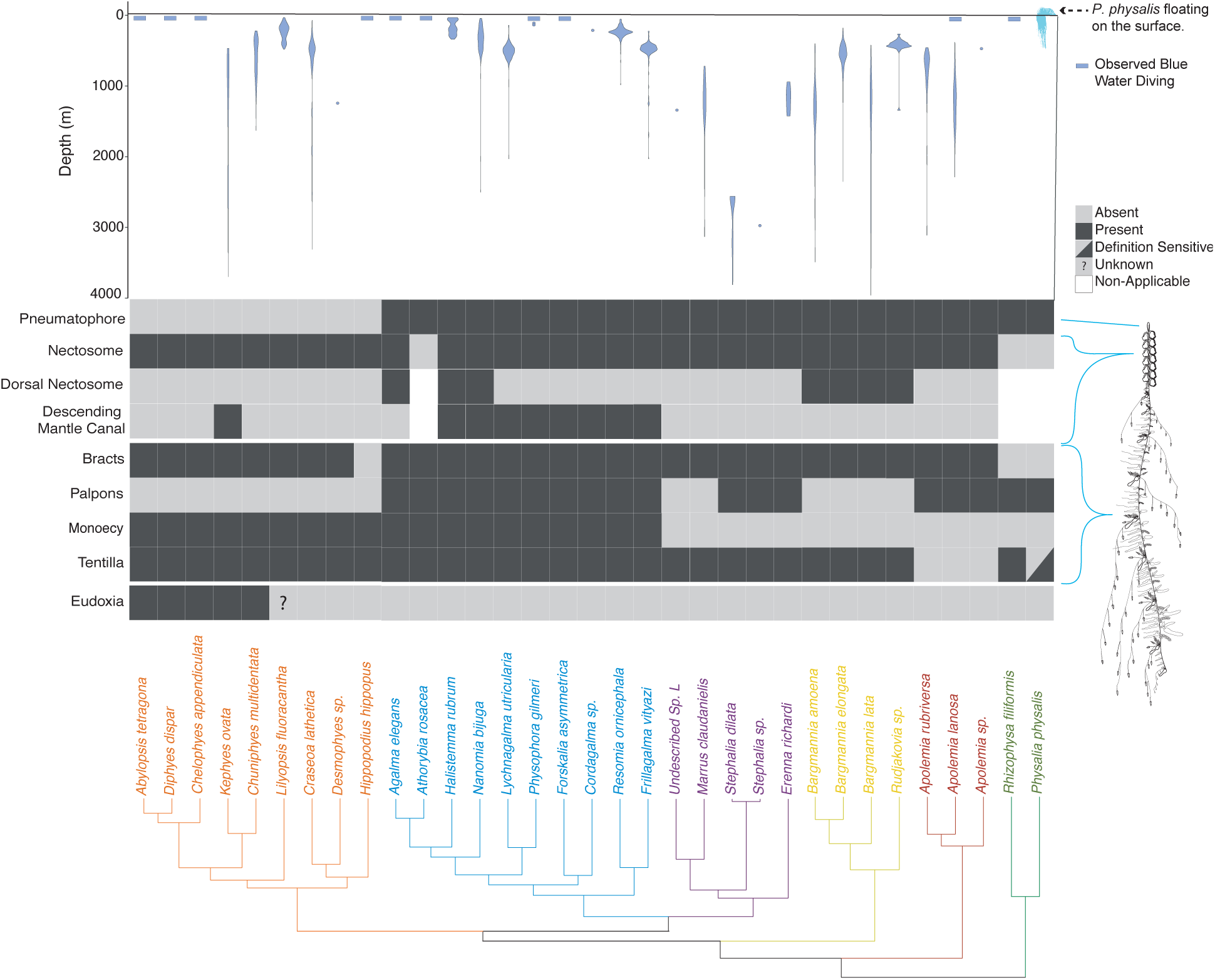
Siphonophore phylogeny showing the distribution of the main anatomical characters and the bathymetric distributions of the different species. Bottom: siphonophore phylogeny, colored by clade. Middle panel: diagram showing the presence/absence of traits across Siphonophora, with the physical location of the trait shown on a schematic of *Nanomia bijuga* (schematic by Freya Goetz). Top: Bathymetric distribution of siphonophore species. Physalia illustration by Noah Schlottman, taken from http://phylopic.org/

### 3.2 Character Evolution

### 3.3 Evolution of Monoecy

In all siphonophores, each gonophore (sexual medusa that produces gametes) is either male or female. Within each siphonophore species, colonies are either monoecious (male and female gonophores are on the same colony) or dioecious (male and female gonophores are on different colonies). Previous analyses suggested that the common ancestor of siphonophores was dioecious, and was consistent with a single gain of monoecy within Codonophora and no secondary losses (Dunn et al., 2005). The better-resolved tree in the current analyses indicates that the evolution of monoecy is more complicated than this. The two clades of monoecious siphonophores, Calycophorae and Clade A (Fig. 3A), do not form a monophyletic group. This is because Clade B, which contains dioecious species, is also descended from their most recent common ancestor. SOWH tests strongly reject the placement of the monecious clades Calycophorae and Clade A as a group that excludes Clade B (Figs. 3C and S2). The positions of the only two taxa from Clade B in the previous analysis (Dunn et al., 2005), *Erenna* and *Stephalia*, were unresolved in that previous study. This difference in conclusions regarding trait evolution, therefore, does not reflect a contradiction between alternative strongly supported results, but the resolution of earlier polytomies in a way that indicates there has been homoplasy in the evolution of monoecy.

The distribution of monoecy is consistent with two scenarios (Fig. 4). In the first, there is a single shift from dioecy to monoecy along the branch that give rise to the most recent common ancestor of Calycophorae and Euphysonectae, followed by a shift back to dioecy along the branch that gave rise to Clade B. In the second, monoecy arose twice – once along the branch that gave rise to Clade A and again along the branch that gave rise to Calycophorae.

Ancestral character state reconstructions favor the hypothesis that monoecy arose twice (Fig. 5A), once in Calycophorae and once in Clade A. This is consistent with differences in the arrangements of male and female gonophores in the two clades. In Clade A, male and female zooids are found within the same cormidium (a single reiterated sequence of zooids along the stem, see Fig. 2). In these species, the male and female zooids are placed at very different but well defined locations within the cormidium. Meanwhile in calycophorans, each cormidium bears either male or female gonophores. In this form of monoecy, the male and female cormidia can either be in an alternating pattern, or there can be several male or female cormidia in a row. In either case, male and female zooids are found at the same corresponding locations within the cormidia. In sum, homoplasy in sexual system evolution along with variation in the spatial arrangement of gonophores within a colony suggest that siphonophores have evolved different ways to be monoecious. Both Calycophorae and Clade A have a large proportion of shallow water species (see section 3.7), suggesting that there may be an association between habitat depth and sexual mode. Similar independent transitions from gonochorism (separate sex) to hermaphroditism (both sexes in the same individual) have been identified in shallow-water scleractinian corals (Anthozoa, Cnidaria) (Kerr et al., 2011). To test this hypothesis, a more extensive taxon sampling of the Calycophorae is needed.

**Figure 5:**
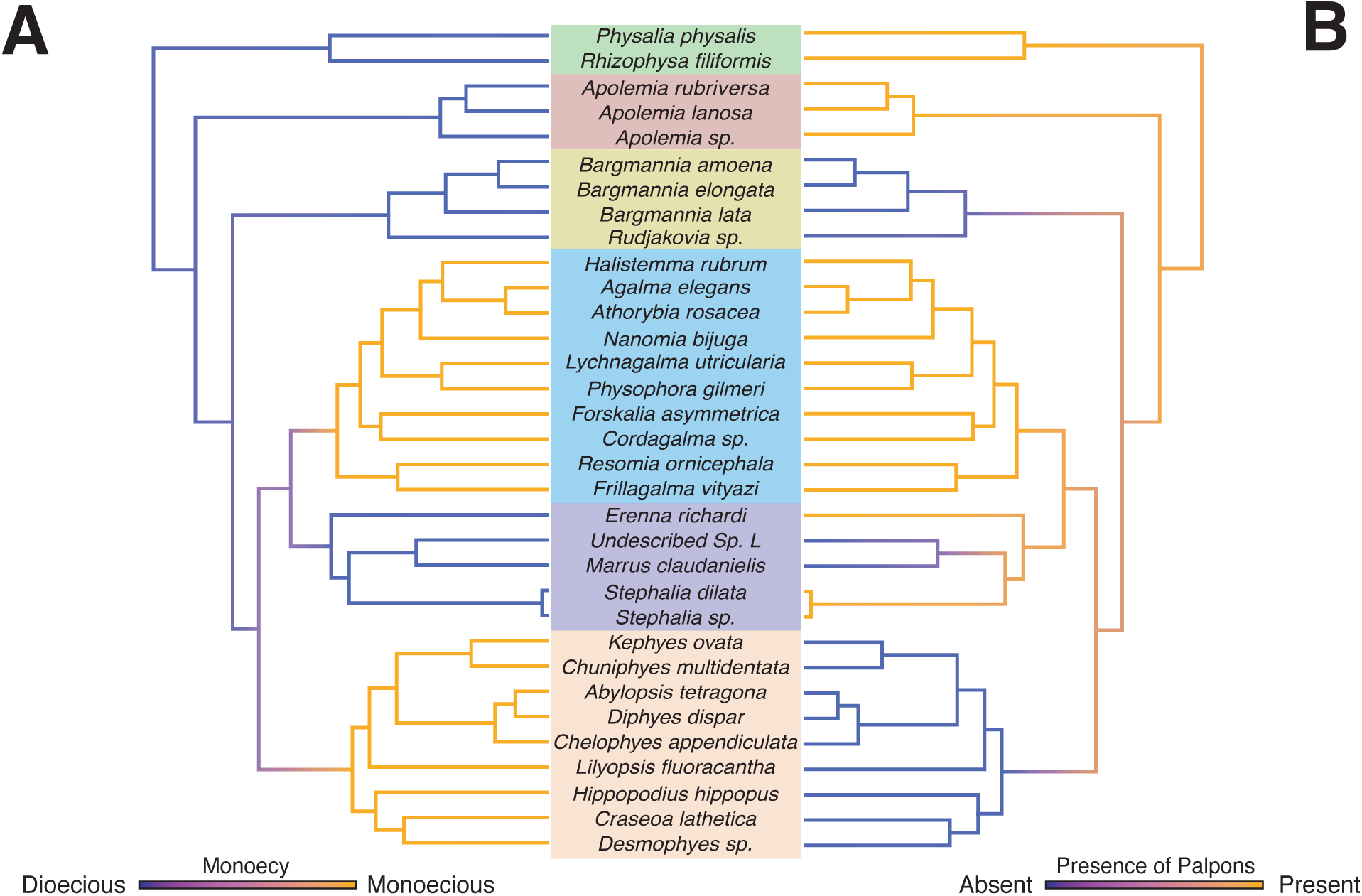
Stochastic mapping reconstruction of the evolutionary history of A) sexual mode, whether a colony is monoecious or dioecious and B) presence/absence of palpons (modified reduced gastrozooids). The color gradients show the reconstructed probability estimate of the discrete character states along the branches. Intermediate values reflect uncertainty.

Within Calycophorae there are some variations in sexual mode – in *Sulculeolaria* (not included in this phylogeny) colonies appear to consist of only one sex, however they are monoecious and protandrous, with female gonophores developing after the release of male gonophores (Carré, 1979). Environmental influences may also play a role in determining the expressed sex. Colonies of the calycophoran *Chelophyes appendiculata* collected in the field always bear both male and female gonophores whereas when kept in culture only gonophores of one sex are maintained (Carré and Carré, 2000). This suggests a high plasticity of the sexual state in some calycophoran taxa and underlines the need for caution when evaluating the state of this character in rarely collected species.

### 3.4 The Evolution of Zooid Types

One of the most striking aspects of siphonophore biology is their diversity of unique zooid types (Beklemishev, 1969; Cartwright and Nawrocki, 2010). For example, members of *Forskalia* have 5 basic zooid types (nectophore, gastrozooid, palpon, bract, and gonophore), and in some species, a total of 9 when considering zooid subtypes (4 types of bract, male & female gonophores)(Pugh, 2003). Here we reconstruct the evolutionary origins of several zooid types on the present transcriptome tree (Fig. 4).

Nectophores (Fig. 2) are non-reproductive propulsive medusae. In Codonophora, the nectophores are localized to a region known as the nectosome (Fig. 2B), which has its own growth zone, and are used for coordinated colony-level swimming. Planktonic cystonects like *Bathyphysa sibogae* and *Rhizophysa filiformis* (Fig. 1A) instead move through the water column using repeated contraction and relaxation of the stem, and use modified flattened gastrozooids with wings (called ptera) to increase surface area and prevent colony sinking (Biggs and Harbison, 1976). Nectophores are also present within the gonodendra (reproductive structures) of cystonects, and are thought to propel the gonodendra when they detach from the colony (Totton, 1960, Totton, 1965). It is not clear whether the nectophores found within the siphosome of the cystonects are homologous to the nectophores borne on the nectosome of codonophorans. Similarly, the homology of the special nectophore associated with gonophores of the calycophoran *Stephanophyes superba* is also unclear (Chun, 1891). As such, we only consider the evolution of the nectosome in this study, and not the presence/absence of nectophores. The present analyses, as well as the analyses of Dunn et al. (2005), are consistent with a single origin of the nectosome (Fig. S5).

Within the nectosome, the nectophores can be attached along the dorsal or ventral side of the stem, following the orientation framework of Haddock et al. (2005). Our ancestral reconstructions for this character (Fig. S7) suggest that ventral attachment of nectophores was the ancestral state in Codonophora, and that dorsal attachment has independently evolved twice – once along the stem of Agalmatidae and once along the stem of Pyrostephidae. The functional implication of dorsal vs. ventral attachment is not clear.

Bracts are highly reduced zooids unique to siphonophores, where they are only present in Codonophora (Fig. 4). Bracts are functional for protection of the delicate zooids and to help maintain neutral buoyancy (Jacobs, 1937). Some calycophorans are able to actively exclude sulphate ions in their bracts to adjust their buoyancy along the colony (Bidigare and Biggs, 1980). Bracts were lost in Hippopodiidae, some clausophyids, *Physophora hydrostatica* (Fig. 1K), and in *Gymnopraia lapislazula*. These patterns of loss are not captured in this study, as most of these species are not included in the present phylogeny. In species without bracts, a diversity of zooids appear to fulfil the roles of neutral buoyancy and protection. In *Physophora hydrostatica*, enlarged palpons surround all other siphosomal zooids and move in a coordinated manner to inflict a powerful sting (Totton, 1965). While *Hippopodius hippopus* have up to twelve nectophores and can retract the stem within the nectophores – the nectophores play a role in maintaining neutral buoyancy and possibly also in defense, by bioluminescing and blanching in response to stimuli (Fig. 1C shows the blanching of nectophores)(Bassot et al., 1978).

Palpons are modified reduced gastrozooids used for digestion and circulation of the gastrovascular fluid (Mackie et al., 1987). We do not distinguish here between gonopalpons (palpons associated with gonodendra, without a tentacle, as in the cystonects) and palpons borne on the stem (typically with a reduced tentacle or palpacle) (Totton, 1965). We reconstruct them as present in the common ancestor of siphonophores (Fig. 5B), retained in most species, but lost three times independently in the branches leading to Pyrostephidae (represented here by the genera *Bargmannia* and *Rudjakovia*), in calycophorans, and in *Marrus claudanielis*. Within the calycophorans, one species *Stephanophyes superba* (not included in this phylogeny) has polyp-like zooids that have been described as palpons (Totton, 1965), but the exact identity of this zooid is not clear and needs further morphological examination.

### 3.5 The Gain and Loss of the Pneumatophore

The pneumatophore is a gas filled float located at the anterior end of the colony, which helps the colony maintain its orientation in the water column, and plays a role in flotation in the case of the cystonects (Samuel H Church et al., 2015; Mackie, 1974; Totton, 1965). It is not a zooid, as it is not formed by budding but by invagination at the aboral end of the planula during early development (Carré, 1969; Garstang, 1946; Leloup, 1935). Recent descriptions of the neural arrangement in the pneumatophore of *Nanomia bijuga* suggests it could also gather information on relative pressure changes (and thus depth changes), helping regulate geotaxis (Samuel H Church et al., 2015). The ancestral siphonophore had a pneumatophore (Fig. 2B), since both cystonects and all “physonects” possess one (Fig. 4). The pneumatophore was lost in Calycophorae and never gained again in that clade. Calycophorans rely on the ionic balance of their gelatinous nectophores and bracts to retain posture and neutral buoyancy (Mackie, 1974).

### 3.6 The Gain and Loss of Tentilla

Gastrozooids (specialized feeding polyps) have a single tentacle attached to the base of the zooid that is used for prey capture (with the exception of *Physalia physalis*, which has separate zooids for feeding and prey capture). As in other cnidarians, nematocyst stinging capsules, arranged in dense nematocyst batteries, play a critical role in prey capture. In many siphonophore species these batteries are found in side branches of the tentacle, termed tentilla (Fig. 2A). Outside of Siphonophora, most hydrozoans bear simple tentacles without side branches. It is still an open question whether the common ancestor of Siphonophora had tentilla. The only siphonophores species regarded as lacking tentilla are *Physalia physalis*, *Apolemia* spp. (Fig. 1H), and *Bathyphysa conifera* (Fig. 1B). Since *B. conifera* is the only member of the *Rhizophysidae* (and of the *Bathyphysa* genus) lacking tentilla, we assume this is a case of secondary loss. When we reconstruct the evolution of this character on the current phylogeny, 70%sss of simulations support a common ancestor bearing tentilla, with two independent losses leading to *Physalia* and *Apolemia* (Fig. S3). However, this leaves a 30%sss support for a simple-tentacled common ancestor followed by 2 independent gains of tentilla in the branches leading to *Rhizophysidae* and non-apolemiid codonophorans.

How we define absence of tentilla, especially for *Physalia physalis*, is also important. The tentacles of this species, when uncoiled, show very prominent, evenly spaced, bulging buttons which contain in the ectoderm all functional nematocytes (carrying mature nematocysts) used by the organism for prey capture (Hessinger and Ford, 1988; Totton, 1960). Siphonophore tentilla are complete diverticular branchings of the tentacle ectoderm, mesoglea, and gastrovascular canal (lined by endoderm). *Physalia*’s buttons enclose individual fluid-filled chambers connected by narrow channels to the tentacular canal, lined by endoderm (Bardi and Marques, 2007). This suggests they are not just ectodermal swellings, but probably reduced tentilla. When we define *Physalia physalis* as tentilla bearing, the results for the character reconstruction lead to a more robust support for a tentilla-bearing common ancestor followed by independent losses of tentilla in the branch leading to Apolemiidae (Fig. S4), and in *Bathyphysa conifera*. The application of phylogenetic methods to the evolution of tentillum morphology would be a crucial step towards understanding the evolution of these structures, and their relationship with the feeding ecology of siphonophores.

### 3.7 The Evolution of Vertical Habitat Use

Siphonophores are abundant predators in the pelagic realm, ranging from the surface (*Physalia physalis*) to bathypelagic depths (Figs. 4 and S8) (Mackie et al., 1987; Mapstone, 2014). The depth distribution of siphonophore populations is not always static, as some species are known to be vertical migrators, although this is within a relatively narrow depth range (<100m) (Pugh, 1984). Some species such as *Nanomia bijuga* exhibit synchronous diel migration patterns (Barham, 1966). Using the present phylogeny, we reconstructed the median depth changes along the phylogeny under a Brownian Motion model (Fig. S9), which had the strongest AICc support (compared to non-phylogenetic distributions, and to Ohrnstein-Uhlenbeck). This model indicates a mesopelagic most recent common ancestor, with several independent transition events to epipelagic and bathypelagic waters. There was only a single transition to benthic lifestyle on the branch of Rhodaliidae, and a single transition to a pleustonic lifestyle on the branch of *Physalia physalis*. There is evidence that habitat depth is conserved within some clades, with the exception of Calycophorae which have diversified across the water column (Figs. S8, S9). Depth appears to be phylogenetically conserved in Euphysonectae after the split between Clade A (shallow living species) and Clade B (deep dwelling species); however several shallow-living species that likely belong in Clade B were not included in this analysis. The present sampling is also not sufficient to capture significant variation in depth distributions between closely related species. Previous studies have shown that many species that are collected at the same locality are found to occupy discrete, largely non-overlapping depth distributions, including between species that are closely related (Pugh, 1974). This suggests that vertical habitat use is more labile than it appears and may be an important mechanism in siphonophore ecology.

This reconstruction (Fig. S8) only included depths recorded using an ROV, thus it excludes many other independent colonizations of the epipelagic habitat. The ROV observations are reliable below 200m, and no quantitative measurements were made on SCUBA dives. Species such as *Hippopodius hippopus*, *Athorybia rosacea*, *Diphyes dispar*, and *Chelophyes appendiculata* are often encountered blue water diving less than 20m from the surface (Fig. 4). We also reconstructed the median depth changes along the phylogeny using median depths of 20m for all species collected by SCUBA diving or via a shallow trawl (Fig. S9), and still find support for a mesopelagic ancestor. It is important to note, however, that *H. hippopus* and *C. appendiculata* were both collected in the bay of Villefrance-sur-mer, France, where an upwelling is known to bring deeper species closer to the surface (Nival et al., 1976).

## 4. Conclusions

Using phylogenomic tools we were able to resolve deep relationships within Siphonophora with strong support. Among other relationships, we identify the clade Euphysonectae as the sister group to Calycophorae. Our results suggest that monoecy arose twice, based both on phylogenetic reconstruction and differences in the way monoecy is realized in different clades. We are unable to fully capture some of the complex patterns of zooid gain and loss within Codonophora, which will require greater taxon sampling and improved morphological understanding of many poorly known species. The improved resolution presented in this study suggests that an important next step in understanding siphonophore evolution will be targeting molecular sampling within Euphysonectae (where we sampled 13 of 62 valid described species that likely belong to the group) and Calycophorae (where we sampled 9 species in a clade of 109 valid described species) to further resolve the internal relationships within these clades.

## Acknowledgements

This work was supported by the National Science Foundation (DEB-1256695 and the Waterman Award). Sequencing at the Brown Genomics Core facility was supported in part by NIH P30RR031153 and NSF EPSCoR EPS-1004057. Data transfer was supported by NSF RII-C2 EPS-1005789. Analyses were conducted with computational resources and services at the Center for Computation and Visualization at Brown University, supported in part by the NSF EPSCoR EPS-1004057 and the State of Rhode Island. SOWHAT analyses were carried out on the Odyssey cluster supported by the FAS Division of Science, Research Computing Group at Harvard University – we thank Cassandra Extavour for use of the Harvard cluster. We thank Rob Sherlock for providing the *Bathyphysa conifera* photograph. We also thank Zack Lewis for help with RNA extractions. We also thank the MBARI crews and ROV pilots for collection of the specimens.

## Supplementary Information

### Agalma analysis

**Figure S1:**
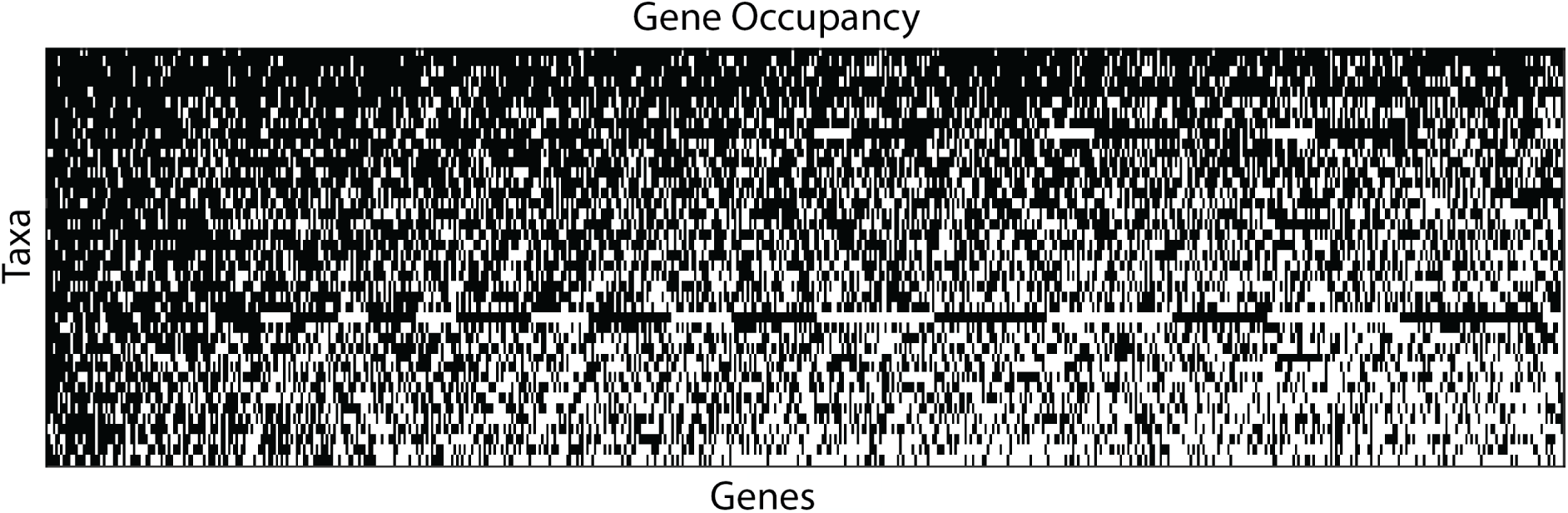
60%sss gene occupancy matrix for 41 species across 1,071 genes. Genes and species are sorted by sampling, the best sampled shown in the upper left.

### SOWH analysis

**Figure S2:**
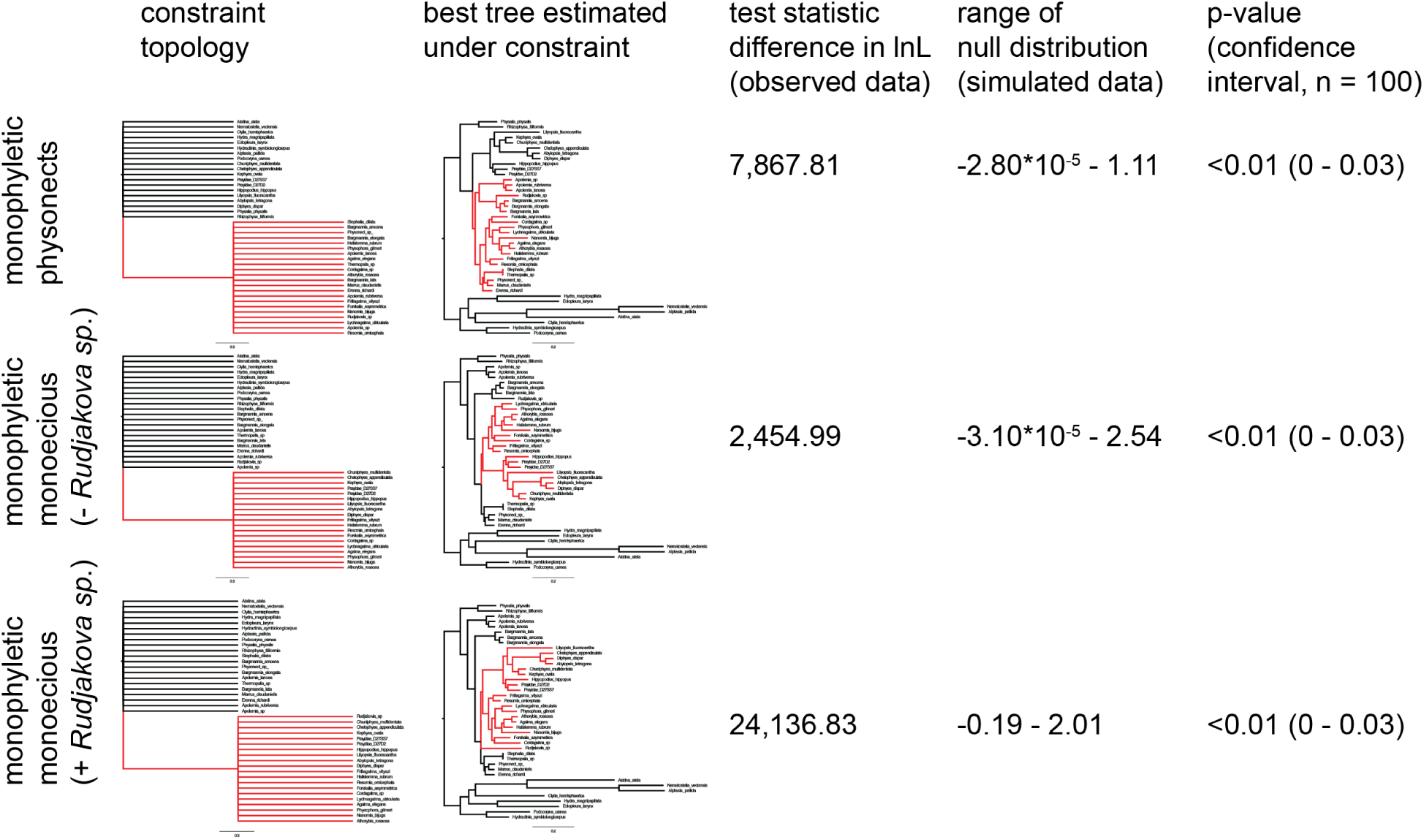
Constrained topologies specified in SOWH testing. Test statistic and p-value for each tree estimated under constraint are given.

### Stochastic Character maps

**Figure S3:**
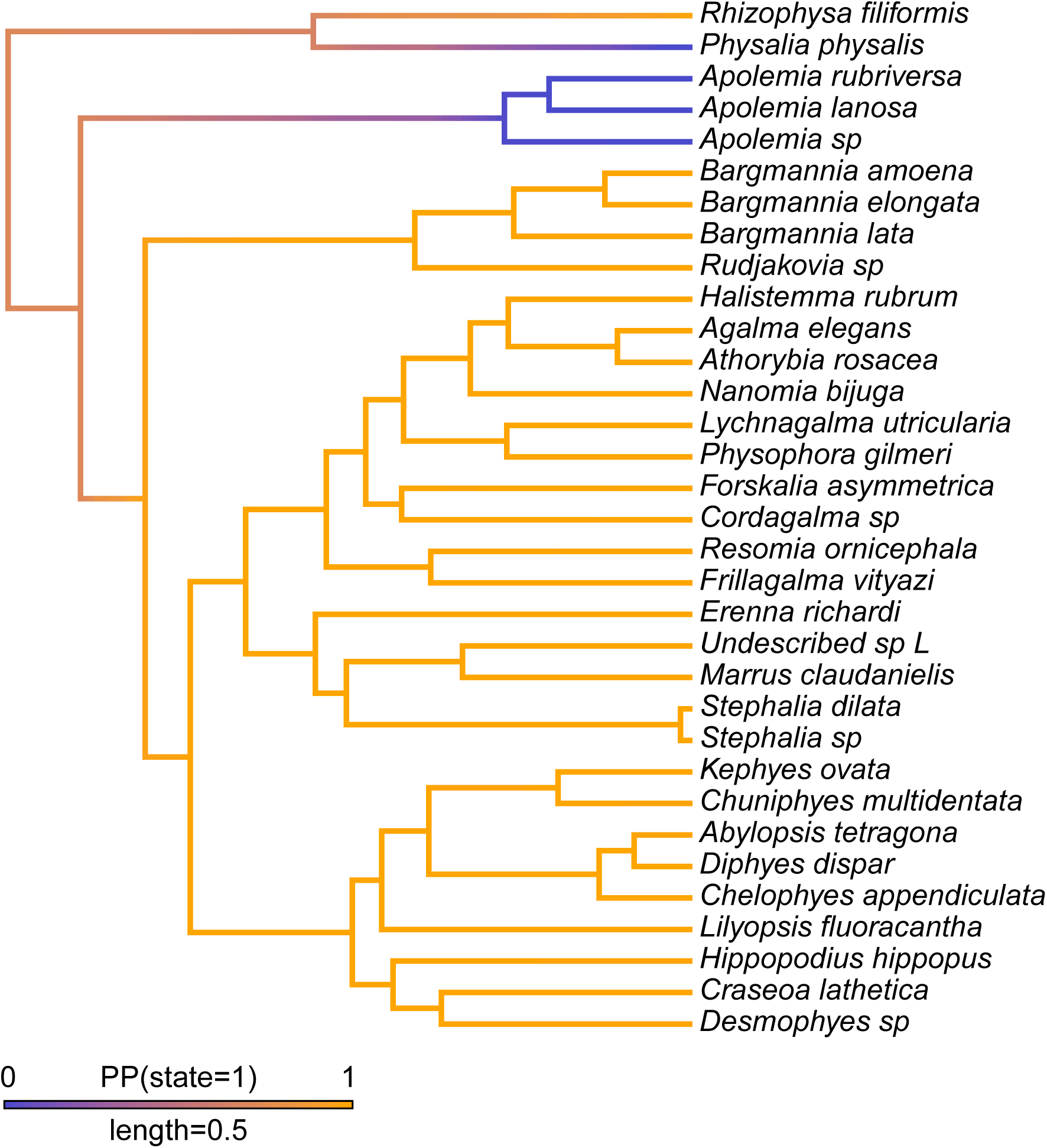
Stochastic character map of presence of tentilla with *Physalia* included as not bearing tentilla.

**Figure S4:**
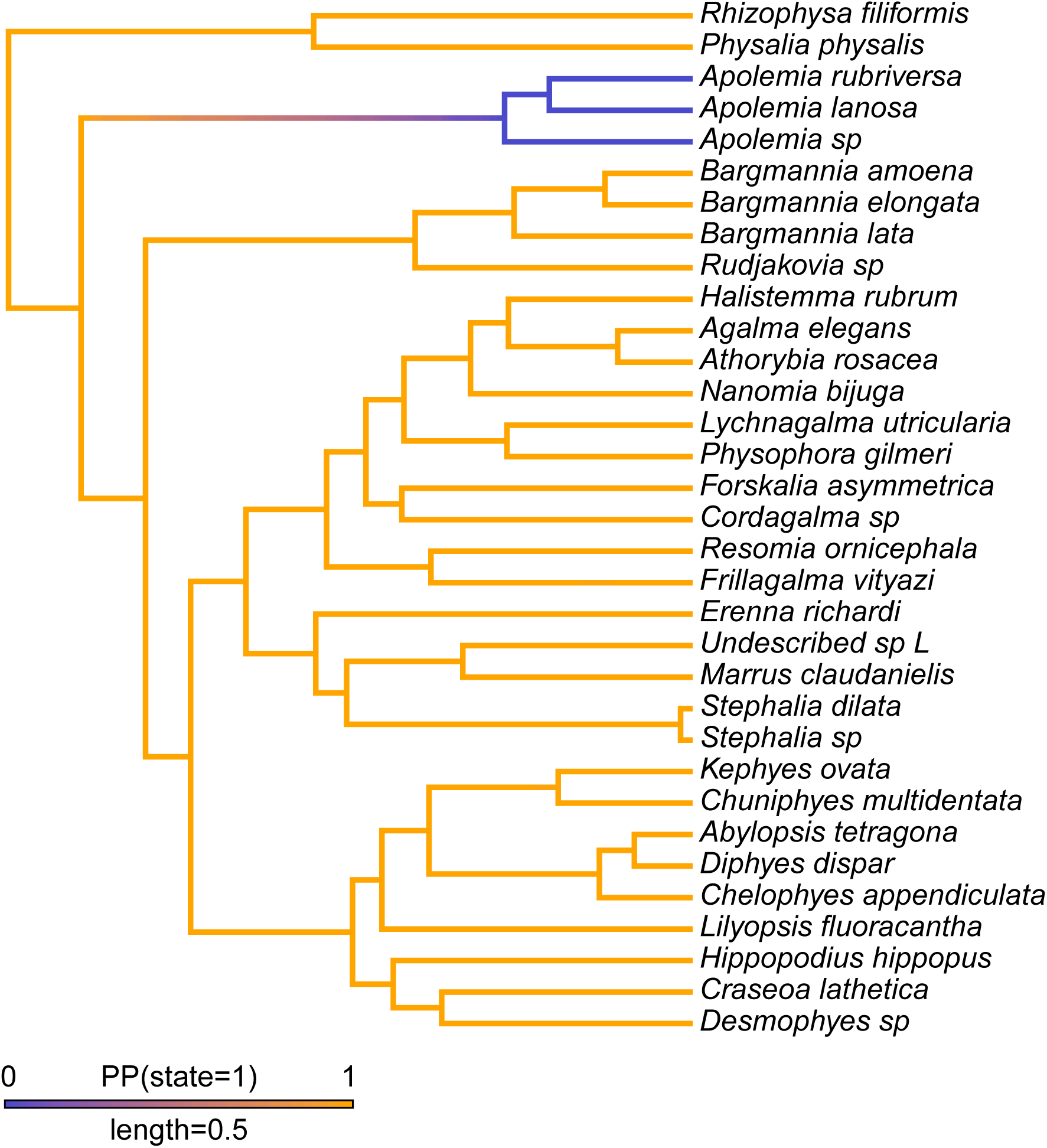
Stochastic character map of presence of tentilla with *Physalia* included as bearing tentilla.

**Figure S5:**
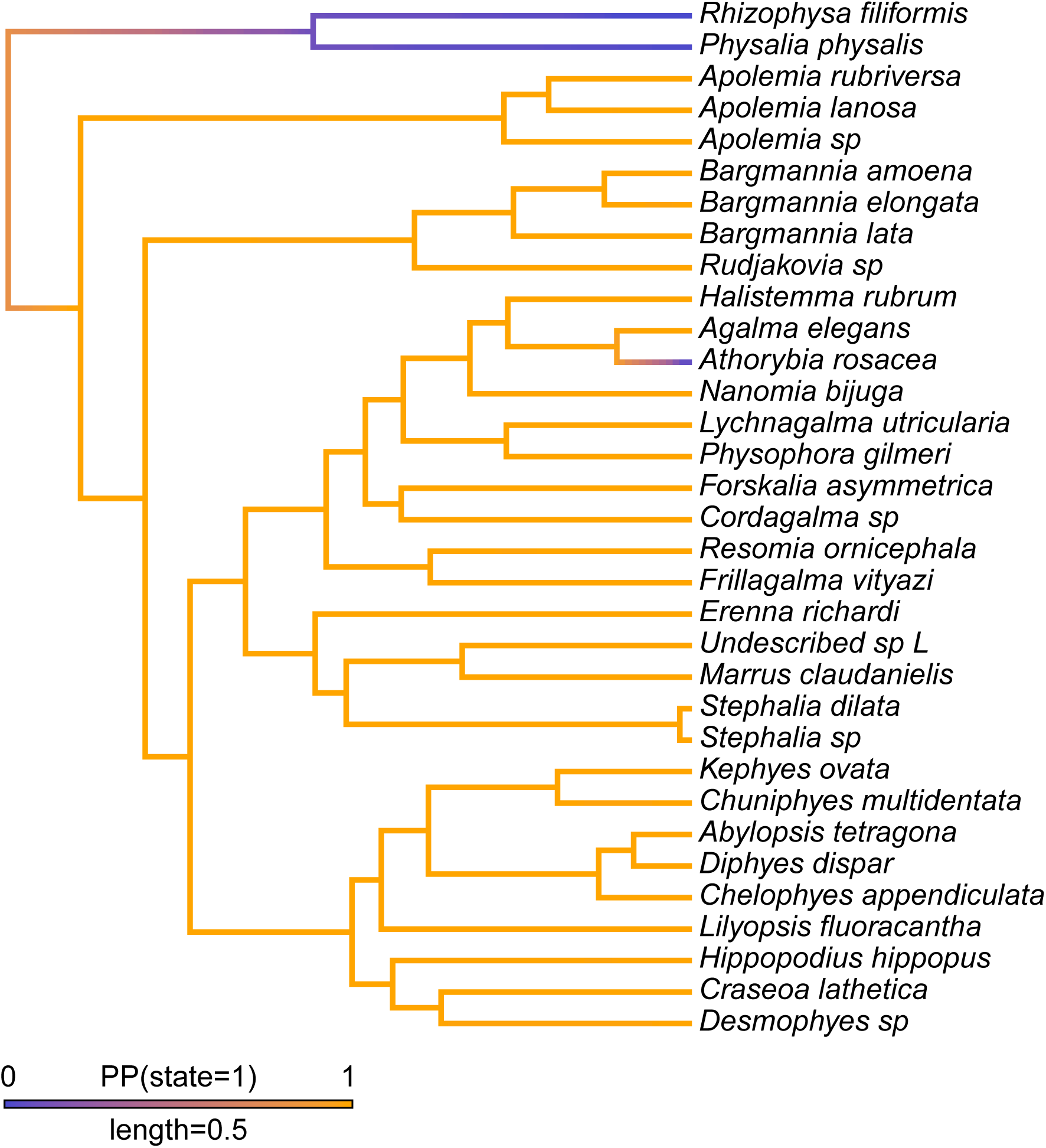
Stochastic character map of presence of nectosome.

**Figure S6:**
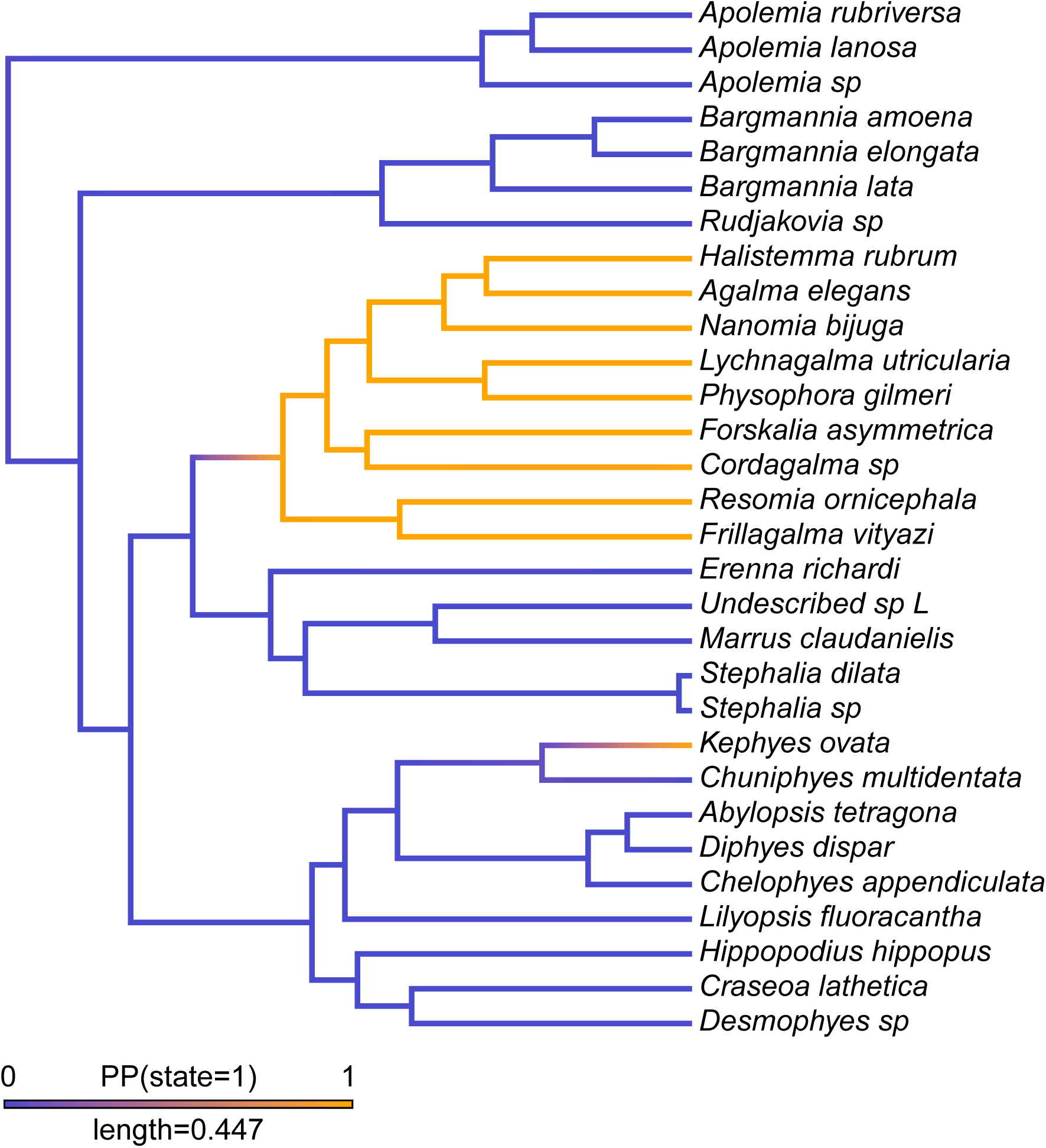
Stochastic character map of presence of a descending mantle canal in the nectophores. Cystonects and Athorybia were excluded as they do not have a nectosome.

**Figure S7:**
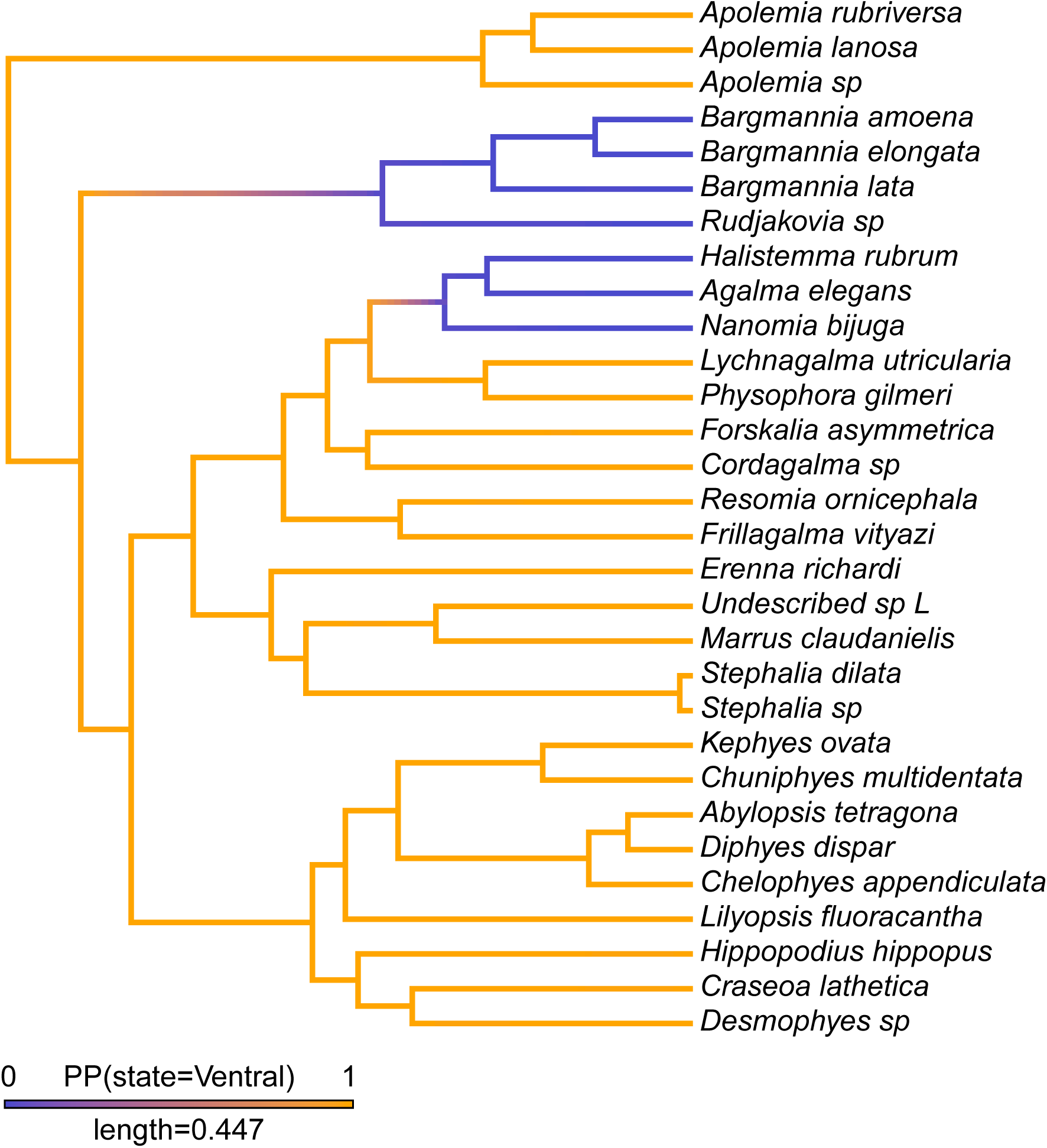
Stochastic character map for the evolution of the position of the nectosome. Cystonects were excluded as they do not have a nectosome.

**Figure S8:**
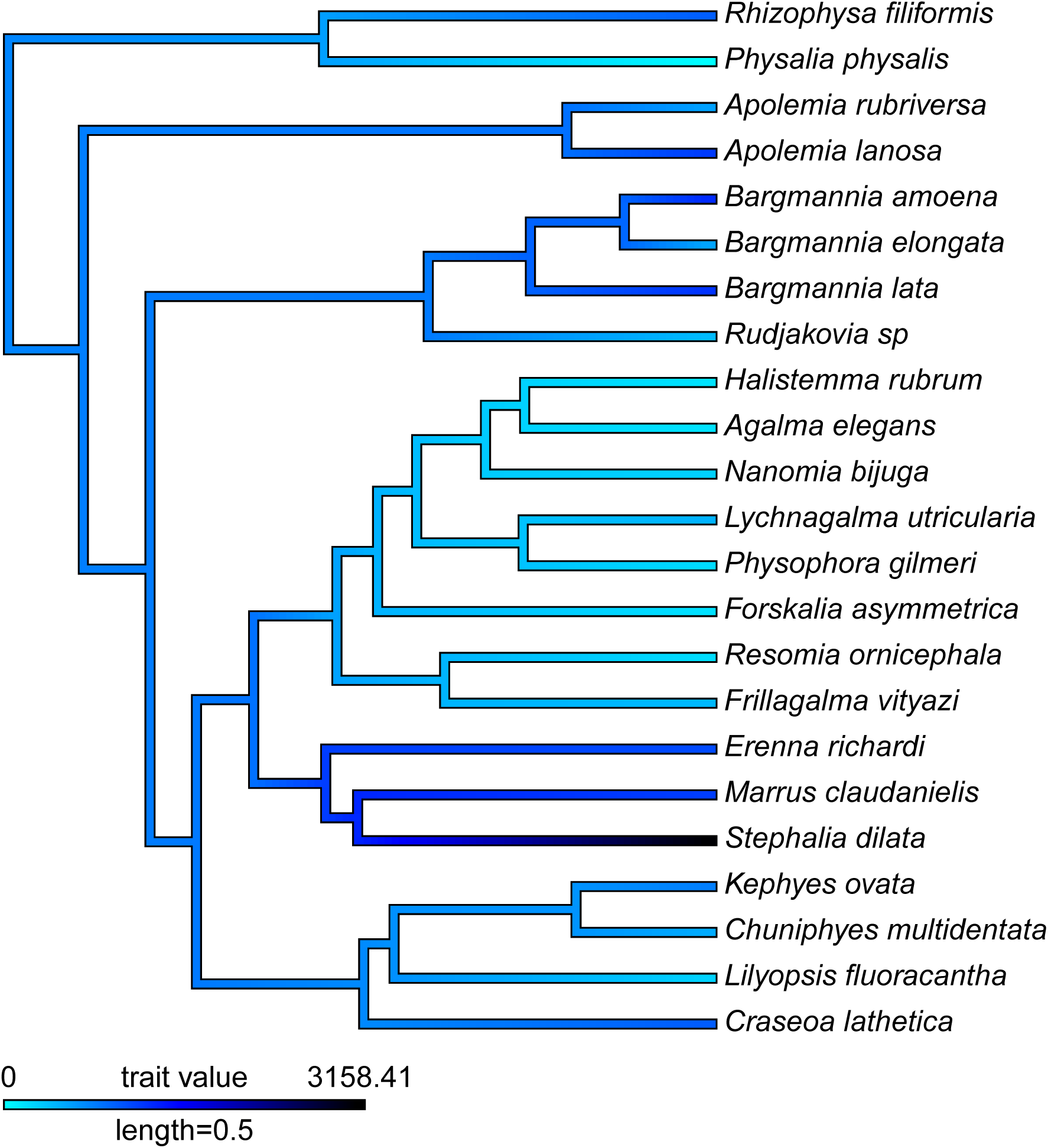
Brownian Motion character map of median depth of species observed with an MBARI ROV.

**Figure S9:**
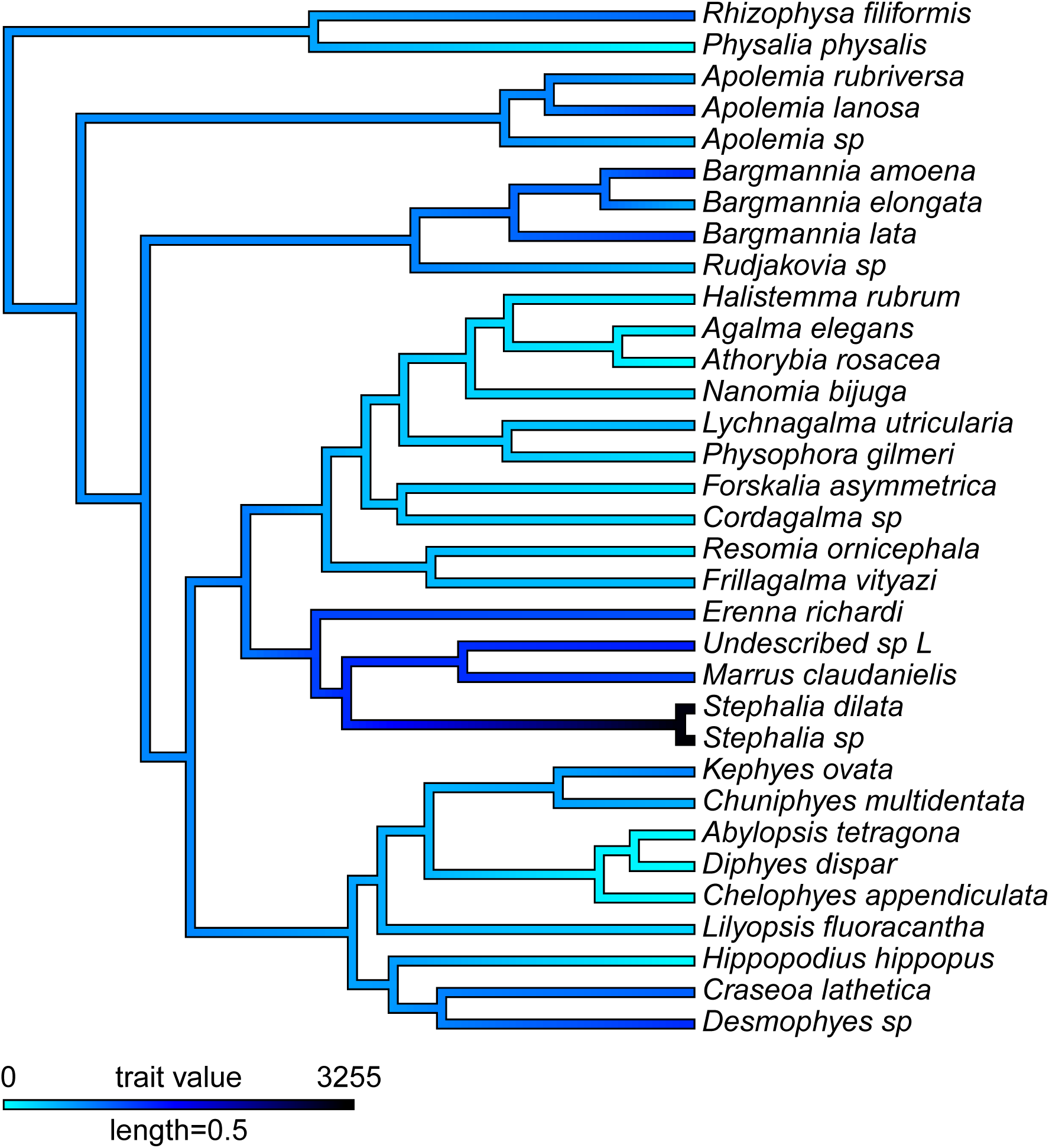
Brownian Motion character map of median depth of species including blue water diving observations.

### Software versions

This manuscript was computed on Fri Jan 19 20:06:20 2018 with the following R package versions.

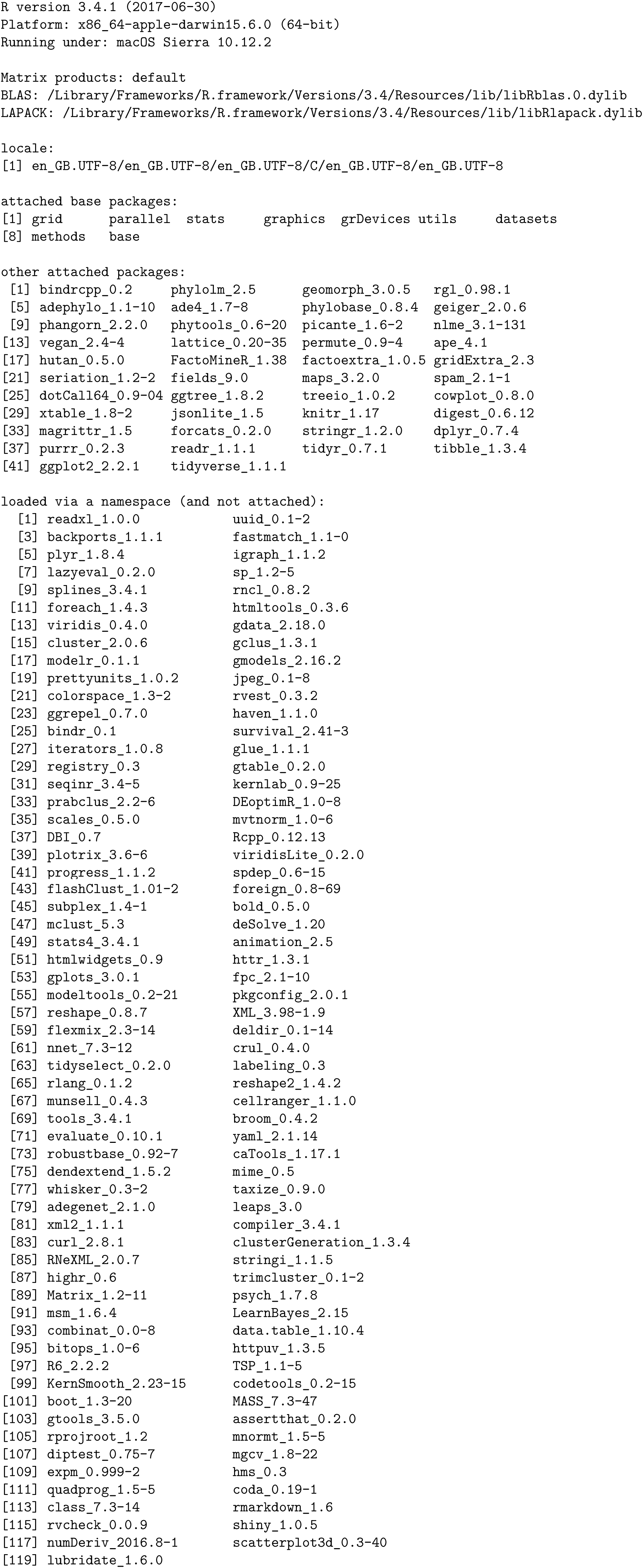

## References

Altschul, S.F., Gish, W., Miller, W., Myers, E.W., Lipman, D.J., 1990. Basic local alignment search tool. J. Mol. Biol. 215, 403–410.

Bardi, J., Marques, A., 2007. Taxonomic redescription of the Portuguese man-of-war, *Physalia physalis* (Cnidaria, Hydrozoa, Siphonophorae, Cystonectae) from Brazil. Iheringia. Ser. Zool. 97, 425–433.

Barham, E.G., 1966. Deep scattering layer migration and composition: observations from a diving saucer. Science 151, 1399–1403.

Bassot, J.-M., Bilbaut, A., Mackie, G., Passano, L., De Ceccatty, M.P., 1978. Bioluminescence and other responses spread by epithelial conduction in the siphonophore Hippopodius. Biol. Bull. 155, 473–498.

Beklemishev, W.N., 1969. Principles of Comparative Anatomy of Invertebrates. Volume I. Promorphology Oliver & Boyd, Edinburgh.

Bidigare, R.R., Biggs, D.C., 1980. The role of sulfate exclusion in buoyancy maintenance by siphonophores and other oceanic gelatinous zooplankton. Comp. Biochem. Physiol. A Physiol. 66, 467–471.

Biggs, D., Harbison, G., 1976. The siphonophore Bathyphysa sibogae Lens and Van Riemsdijk, 1908, in the Sargasso Sea, with notes on its natural history. Bull. Mar. Sci. 26, 14–18.

Carré, C., 1979. Sur le genre Sulculeolaria Blainville, 1834 (Siphonophora, Calycophorae, Diphyidae). Ann. Inst. Oceanogr. (Paris) 55, 27–48.

Carré, C., Carré, D., 1995. Ordre des siphonophores, in: Grassé, P.-P. (Ed.), Traité de Zoologie. Anatomie, Systématique, Biologie. Paris:Masson, pp. 523–596.

Carré, C., Carré, D., 1991. A complete life cycle of the calycophoran siphonophore Muggiaea kochi (Will) in the laboratory, under different temperature conditions: ecological implications. Philos. Trans. R. Soc. Lond., B, Biol. Sci. 334, 27–32.

Carré, D., 1969. Etude histologique du developpement de Nanomia bijuga (Chiaje, 1841), Siphonophore Physonecte, Agalmidae. Cah. Biol. Mar. 10, 325–341.

Carré, D., Carré, C., 2000. Origin of germ cells, sex determination, and sex inversion in medusae of the genus Clytia (Hydrozoa, Leptomedusae): The influence of temperature. J. Exp. Zool. 287, 233–242.

Carré, C, 1967. Le developpement larvaire d’*Abylopsis tetragona*. Cah. Biol. Mar. 8, 185–193.

Cartwright, P., Evans, N.M., Dunn, C.W., Marques, A., Miglietta, M.P., Schuchert, P., Collins, A.G., 2008. Phylogenetics of Hydroidolina (Hydrozoa: Cnidaria). J. Mar. Biol. Assoc. U.K. 88, 1663. https://doi.org/10.1017/S0025315408002257

Cartwright, P., Nawrocki, A.M., 2010. Character evolution in Hydrozoa (phylum Cnidaria). Integr. Comp. Biol. 50, 456–472. https://doi.org/https://doi.org/10.1093/icb/icq089

Choy, C.A., Haddock, S.H.D., Robison, B.H., 2017. Deep pelagic food web structure as revealed by in situ feeding observations. Proc. R. Soc. Lond., B, Biol. Sci. 284. https://doi.org/10.1098/rspb.2017.2017.2116

Chun, C., 1891. Die Canarischen Siphonophoren I. *Stephanophyes superba* und die Familie der Stephanophyi-iden. Abh. Senckenb. Naturforsch. Ges. 16, 553–627.

Church, S.H., Ryan, J.F., Dunn, C.W., 2015. Automation and Evaluation of the SOWH Test with SOWHAT. Syst. Biol. 64, 1048–1058. https://doi.org/10.1093/sysbio/syv055

Church, S.H., Siebert, S., Bhattacharyya, P., Dunn, C.W., 2015. The histology of *Nanomia bijuga* (Hydrozoa: Siphonophora). J. Exp. Zool. B Mol. Dev. Evol. 324, 435–449.

Dunn, C., Pugh, P., Haddock, S., 2005. Molecular Phylogenetics of the Siphonophora (Cnidaria), with Implications for the Evolution of Functional Specialization. Syst. Biol. 54, 916–935. https://doi.org/10.1080/10635150500354837

Dunn, C.W., Howison, M., Zapata, F., 2013. Agalma: an automated phylogenomics workflow. BMC Bioinformatics 14, 330. https://doi.org/10.1186/1471-2105-14-330

Dunn, C.W., Wagner, G.P., 2006. The evolution of colony-level development in the Siphonophora (Cnidaria: Hydrozoa). Dev. Genes Evol. 216, 743–754.

Enright, A.J., Van Dongen, S., Ouzounis, C.A., 2002. An efficient algorithm for large-scale detection of protein families. Nucleic Acids Res. 30, 1575–1584. https://doi.org/10.1093/nar/30.7.1575

Garstang, W., 1946. The morphology and relations of the Siphonophora. Q.J. Microsc. Sci 87, 103–193.

Grabherr, M.G., Haas, B.J., Yassour, M., Levin, J.Z., Thompson, D.A., Amit, I., Adiconis, X., Fan, L., Raychowdhury, R., Zeng, Q., others, 2011. Full-length transcriptome assembly from RNA-Seq data without a reference genome. Nat. Biotechnol. 29, 644–652.

Guang, A., Howison, M., Zapata, F., Lawrence, C.E., Dunn, C., 2017. Revising transcriptome assemblies with phylogenetic information in Agalma 1.0. bioRxiv. https://doi.org/10.1101/202416

Haddock, S.H., Dunn, C.W., Pugh, P.R., 2005. A re-examination of siphonophore terminology and morphology, applied to the description of two new prayine species with remarkable bio-optical properties. J. Mar. Biol. Assoc. U.K. 85, 695–707. https://doi.org/10.1017/S0025315405011616

Hessinger, D., Ford, H., 1988. Ultrastructure of the small cnidocyte of the Portuguese man-of-war (*Physalia physalis*) tentacle, in: Hessinger, D., Lenhoff, H. (Eds.), The Biology of Nematocysts. Academic Press, San Diego, USA.

Huelsenbeck, J.P., Nielsen, R., Bollback, J.P., 2003. Stochastic mapping of morphological characters. Syst. Biol. 52, 131–158.

Jacobs, W., 1937. Beobachtungen Über das Schweben der Siphonophoren. J. Comp. Physiol. A Neuroethol. Sens. Neural. Behav. Physiol. 24, 583–601

Katoh, K., Standley, D.M., 2013. MAFFT multiple sequence alignment software version 7: improvements in performance and usability. Mol. Biol. Evol. 30, 772–780.

Kerr, A.M., Baird, A.H., Hughes, T.P., 2011. Correlated evolution of sex and reproductive mode in corals (anthozoa: Scleractinia). Proc. R. Soc. Lond., B, Biol. Sci. 278, 75–81.

Langmead, B., Salzberg, S.L., 2012. Fast gapped-read alignment with Bowtie 2. Nat. Methods 9, 357–359.

Lartillot, N., Lepage, T., Blanquart, S., 2009. PhyloBayes 3: a Bayesian software package for phylogenetic reconstruction and molecular dating. Bioinformatics25, 2286–2288.

Lartillot, N., Philippe, H., 2004. A bayesian mixture model for across-site heterogeneities in the amino-acid replacement process. Mol. Biol. Evol. 21, 1095–1109. https://doi.org/10.1093/molbev/msh112

Leloup, E., 1935. Les siphonophores de la rade de Villefranche-sur-Mer (Alpes Maritimes, France). Mem. Mus. r. His. nat. Belg. 11, 1–12.

Li, B., Dewey, C.N., 2011. RSEM: accurate transcript quantification from RNA-Seq data with or without a reference genome. BMC Bioinformatics 12, 323.

Li, H., Handsaker, B., Wysoker, A., Fennell, T., Ruan, J., Homer, N., Marth, G., Abecasis, G., Durbin, R., 2009. The sequence alignment/map format and SAMtools. Bioinformatics 25, 2078–2079.

Mackie, G., 1974. Locomotion, flotation, and dispersal. Academic Press, New York, USA.

Mackie, G., Pugh, P., Purcell, J., 1987. Siphonophore biology. Adv. Mar. Biol. 24, 97–262.

Mapstone, G.M., 2014. Global diversity and review of Siphonophorae (Cnidaria: Hydrozoa). PLoS One 9, e87737.

Nival, P., Gostan, J., Malara, G., Chara, R., 1976. Evolution du plancton dans la baie de Villefranche-sur-Mer ala fin du printemps (mai et juin 1971), 2. Biomasse de phytoplancton et production primarie. Vie Milieu 26, 47–76.

Pagès, F., Gonzàlez, H., Ramòn, M., Sobarzo, M., Gili, J.-M., 2001. Gelatinous zooplankton assemblages associated with water masses in the Humboldt Current System, and potential predatory impact by Bassia bassensis (Siphonophora: Calycophorae). Mar. Ecol. Prog. Ser. 210, 13–24.

Pugh, P., 2016. A synopsis of the Family Cordagalmatidae fam. nov.(Cnidaria, Siphonophora, Physonectae). Zootaxa 4095, 1–64.

Pugh, P., 2006. The taxonomic status of the genus Moseria (Siphonophora, Physonectae). Zootaxa1343, 1–42.

Pugh, P., 2003. A revision of the family Forskaliidae (Siphonophora, Physonectae). J. Nat. Hist. 37,1281–1327.

Pugh, P., 1984. The diel migrations and distributions within a mesopelagic community in the north east Atlantic. 7. Siphonophores. Prog. Oceanogr. 13, 461–489.

Pugh, P., 1983. Benthic Siphonophores: A Review of the Family Rhodaliidae (Siphonophora, Physonectae). Philos. Trans. R. Soc. Lond., B, Biol. Sci. 301, 165–300.

Pugh, P., 1974. The vertical distribution of the siphonophores collected during the SOND cruise, 1965. J. Mar. Biol. Assoc. U.K. 54, 25–90.

Pugh, P., Pages, F., Boorman, B., 1997. Vertical distribution and abundance of pelagic cnidarians in the eastern Weddell Sea, Antarctica. J. Mar. Biol. Assoc. U.K. 77, 341–360.

Purcell, J., 1981. Dietary composition and diel feeding patterns of epipelagic siphonophores. Mar. Biol. 65, 83–90.

Revell, L.J., 2012. phytools: an R package for phylogenetic comparative biology (and other things). Methods Ecol. Evol. 3, 217–223. https://doi.org/10.1111/j.2041-210X.2011.00169.x

Schlining, B., Stout, N.J., 2006. MBARI’s video annotation and reference system, in: OCEANS 2006. IEEE, pp. 1–5.

Stamatakis, A., 2006. RAxML-VI-HPC: maximum likelihood-based phylogenetic analyses with thousands of taxa and mixed models. Bioinformatics 22, 2688–2690.

Sukumaran, J., Holder, M.T., 2010. DendroPy: a Python library for phylogenetic computing. Bioinformatics 26, 1569–1571.

Swofford, D., Olsen, G., Waddell, P., Hillis, D., 1996. Molecular systematics. 2nd ed. Sunderland (MA): Sinauer Associates.

Talavera, G., Castresana, J., 2007. Improvement of phylogenies after removing divergent and ambiguously aligned blocks from protein sequence alignments. Syst. Biol. 56, 564–577.

Totton, A., 1960. Studies on *Physalia physalis* (L.). Part 1. Natural history and morphology. Discovery Reports 30, 301–368.

Totton, A.K., 1965. A synopsis of the Siphonophora. British Museum (Natural History).

Williams, R., Conway, D., 1981. Vertical distribution and seasonal abundance of Aglantha digitale (OF Müller)(Coelenterata: Trachymedusae) and other planktonic coelenterates in the northeast Atlantic Ocean. J. Plankton Res. 3, 633–643.

Yu, G., Smith, D.K., Zhu, H., Guan, Y., Lam, T.T.-Y., 2016. ggtree: an R package for visualization and annotation of phylogenetic trees with their covariates and other associated data. Methods Ecol. Evol. 8, 28–36. https://doi.org/10.1111/2041-210X.12628

Zapata, F., Goetz, F.E., Smith, S.A., Howison, M., Siebert, S., Church, S.H., Sanders, S.M., Ames, C.L., McFadden, C.S., France, S.C., others, 2015. Phylogenomic analyses support traditional relationships within Cnidaria. PLoS One10, e0139068.

